# Capturing and Selecting Senescence Variation in Wheat

**DOI:** 10.1101/2020.12.06.413641

**Authors:** Elizabeth A. Chapman, Simon Orford, Jacob Lage, Simon Griffiths

## Abstract

Senescence is a highly quantitative trait, but in wheat the genetics underpinning senescence regulation remain relatively unknown. To select senescence variation, and ultimately identify novel genetic regulators, accurate characterisation of senescence phenotypes is essential. When investigating senescence, phenotyping efforts often focus on, or are limited to, visual assessment of the flag leaves. However, senescence is a whole plant process, involving remobilisation and translocation of resources into the developing grain. Furthermore, the temporal progression of senescence poses challenges regarding trait quantification and description, whereupon the different models and approaches applied result in varying definitions of apparently similar metrics.

To gain a holistic understanding of senescence we phenotyped flag leaf and peduncle senescence progression, alongside grain maturation. Reviewing the literature, we identified techniques commonly applied in quantification of senescence variation and developed simple methods to calculate descriptive and discriminatory metrics. To capture senescence dynamism, we developed the idea of calculating thermal time to different flag leaf senescence scores, for which between year Spearman’s rank correlations of *r* ≥ 0.59, *P* < 4.7 × 10^−5^(TT70), identify as an accurate phenotyping method. Following our experience of senescence trait genetic mapping, we recognised the need for singular metrics capable of discriminating senescence variation, identifying Thermal Time to Flag Leaf Senescence score of 70 (TT70) and Mean Peduncle senescence (MeanPed) scores as most informative. Moreover, grain maturity assessments confirmed a previous association between our staygreen traits and grain fill extension, illustrating trait functionality.

Here we review different senescence phenotyping approaches and share our experiences of phenotyping two independent RIL populations segregating for staygreen traits. Together, we direct readers towards senescence phenotyping methods we found most effective, encouraging their use when investigating and discriminating senescence variation of differing genetic bases, and to aid trait selection and weighting in breeding and research programs alike.

## Introduction

Monocarpic senescence is the final stage in wheat development, during which 80 % of leaf nitrogen and phosphorus are re-assimilated into the developing grain (Buchanan-Wollaston, 2007). Senescence is subject to strong environmental and genetic regulation, and prior to visual yellowing and chlorosis up to 50 % of leaf chlorophyll may be lost (Buchanan-Wollaston *et al*., 2005; Borrill *et al*., 2019). Despite this, senescence progression is typically monitored through recording changes in leaf greenness or chlorophyll content over time, either at the individual flag leaf or canopy level (Figure 1, Table 1; Pask *et al*., 2012; Shrestha *et al*., 2012). With reference to CIMMYT crop ontology (Shrestha *et al*., 2012), ‘Flag Leaf Senescence’ (CO_321:0000194) is commonly assessed using a scale from 0 (0 % senescence) to 10 (100 % senescence) (CO_321:0000382; Pask *et al*., 2012). Alternatively, Normalised Differential Vegetation Index (NDVI; CO_321:0000301) or Green NDVI (GNVDI; CO_321:0000961) can be measured using spectral reflectance, where the change in canopy greenness or photosynthetic size provides a more objective measure of senescence (Table 1).

**Figure 1.**
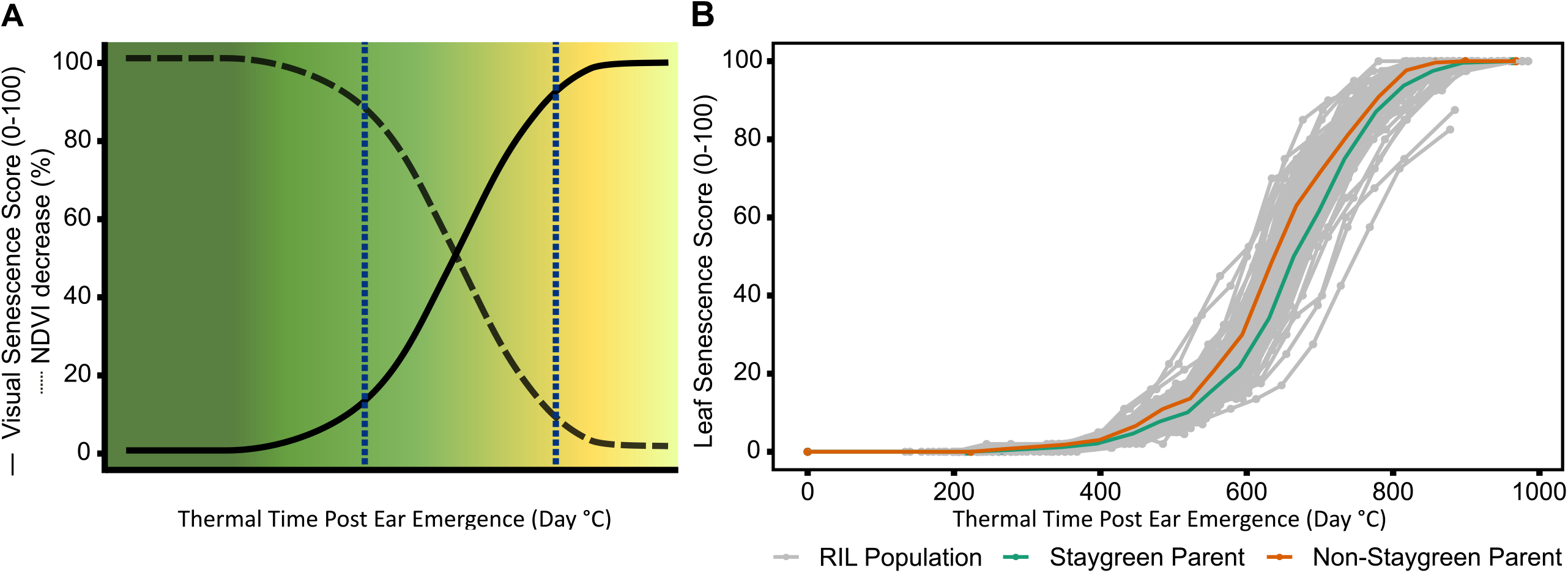
During the rapid senescence phase flag leaves transition from green to yellow (A), with inter-line variation sometimes difficult to distinguish **(B)**. Senescence is scored on a 0-100 scale based in progression of leaf yellowing (visual) or greenness reduction (NDVI-based). Scores are plotted against thermal time post anthesis (^°^ C day) to standardise for heading-date variation. **(B)** Senescence variation recorded for a segregating RIL population (grey, n = 75), Staygreen parent (green), non-staygreen parent (orange); mean visual senescence score (n ≥ 2), Church Farm, Norwich, 2018.

**Table 1.**
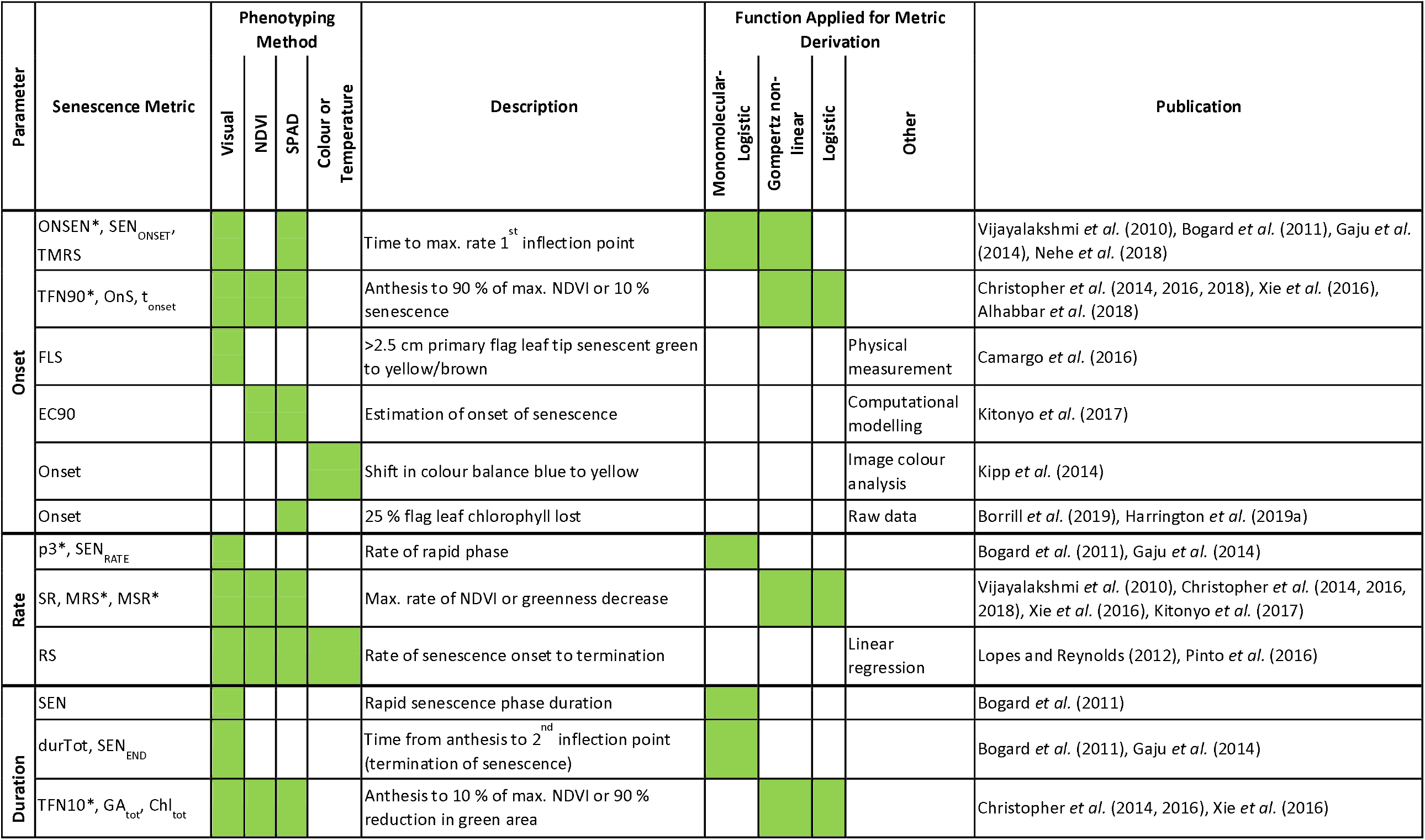
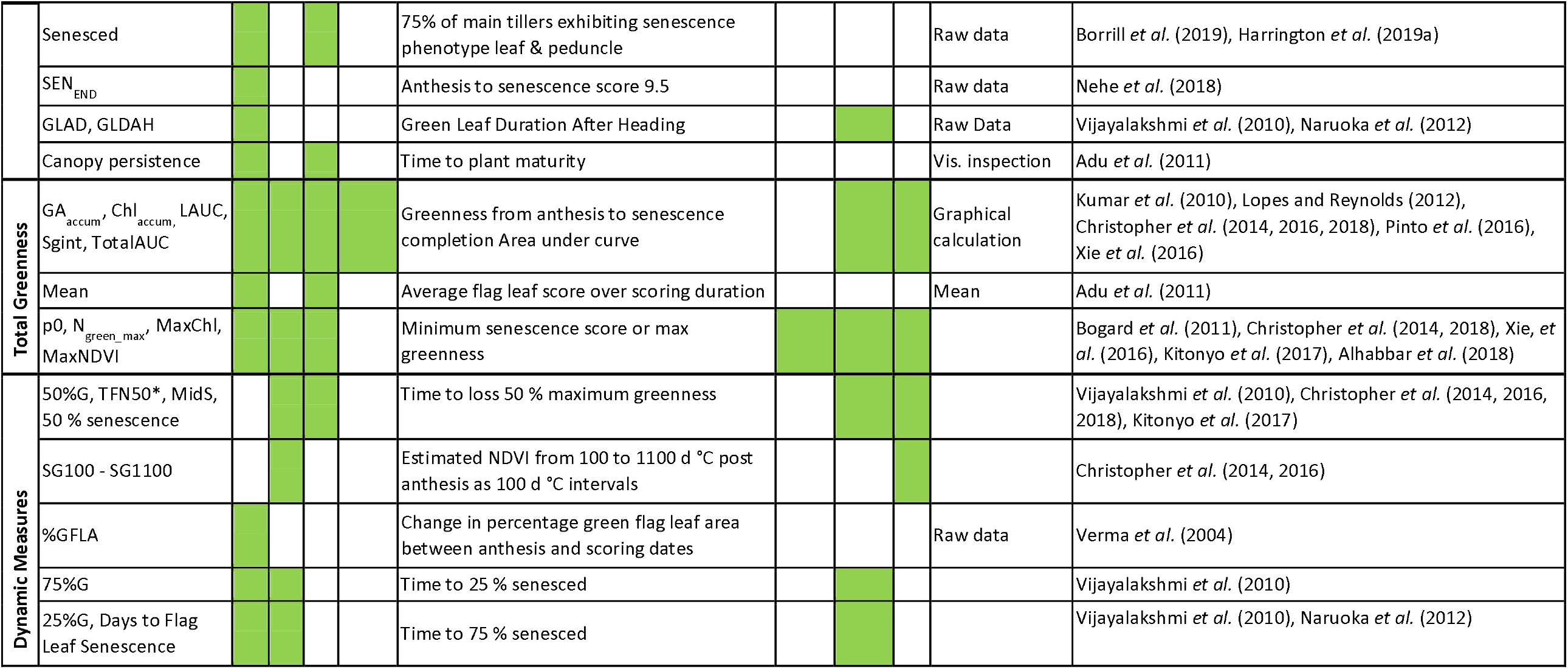
Reviewing methods used to score and quantify leaf senescence. Time-course senescence data can be reduced to a series of parameters using different methods, promoting characterisation and comparison of senescence phenotypes. Shading indicates method of phenotyping and metric calculation.

| Parameter | Senescence Metric | Phenotyping Method |  |  |  | Description | Function Applied for Metric Derivation |  |  |  | Publication |
| --- | --- | --- | --- | --- | --- | --- | --- | --- | --- | --- | --- |
|  |  | Visual | NDVI | SPAD | Colour or Temperature |  | Monomolecular-Logistic | Gompertz non-linear | Logistic | Other |  |
| Onset | ONSEN*, SEN<sub>ONSET'</sub><br>TMRS |  |  |  |  | Time to max. rate 1<sup>st</sup> inflection point |  |  |  |  | Vijayalakshmi *et al.* (2010), Bogard *et al.* (2011), Gaju *et al.* (2014), Nehe *et al.* (2018) |
|  | TFN90*, OnS, t<sub>onset</sub> |  |  |  |  | Anthesis to 90 % of max. NDVI or 10 % senescence |  |  |  |  | Christopher *et al.* (2014, 2016, 2018), Xie *et al.* (2016), Alhabbar *et al.* (2018) |
|  | FLS |  |  |  |  | >2.5 cm primary flag leaf tip senescent green to yellow/brown |  |  |  | Physical measurement | Camargo *et al.* (2016) |
|  | EC90 |  |  |  |  | Estimation of onset of senescence |  |  |  | Computational modelling | Kitonyo *et al.* (2017) |
|  | Onset |  |  |  |  | Shift in colour balance blue to yellow |  |  |  | Image colour analysis | Kipp *et al.* (2014) |
|  | Onset |  |  |  |  | 25 % flag leaf chlorophyll lost |  |  |  | Raw data | Borrill *et al.* (2019), Harrington *et al.* (2019a) |
| Rate | p3*, SEN<sub>RATE</sub> |  |  |  |  | Rate of rapid phase |  |  |  |  | Bogard *et al.* (2011), Gaju *et al.* (2014) |
|  | SR, MRS*, MSR* |  |  |  |  | Max. rate of NDVI or greenness decrease |  |  |  |  | Vijayalakshmi *et al.* (2010), Christopher *et al.* (2014, 2016, 2018), Xie *et al.* (2016), Kitonyo *et al.* (2017) |
|  | RS |  |  |  |  | Rate of senescence onset to termination |  |  |  | Linear regression | Lopes and Reynolds (2012), Pinto *et al.* (2016) |
| Duration | SEN |  |  |  |  | Rapid senescence phase duration |  |  |  |  | Bogard *et al.* (2011) |
|  | durTot, SEN<sub>END</sub> |  |  |  |  | Time from anthesis to 2<sup>nd</sup> inflection point (termination of senescence) |  |  |  |  | Bogard *et al.* (2011), Gaju *et al.* (2014) |
|  | TFN10*, GA<sub>tot'</sub>, Chl<sub>tot</sub> |  |  |  |  | Anthesis to 10 % of max. NDVI or 90 % reduction in green area |  |  |  |  | Christopher *et al.* (2014, 2016), Xie *et al.* (2016) |

|  |  |  |  |  |  |  |  |  |  |  |
| --- | --- | --- | --- | --- | --- | --- | --- | --- | --- | --- |
|  | Senesced |  |  |  | 75% of main tillers exhibiting senescence phenotype leaf & peduncle |  |  |  | Raw data | Borrill *et al.* (2019), Harrington *et al.* (2019a) |
|  | SEN<sub>END</sub> |  |  |  | Anthesis to senescence score 9.5 |  |  |  | Raw data | Nehe *et al.* (2018) |
|  | GLAD, GLDAH |  |  |  | Green Leaf Duration After Heading |  |  |  | Raw Data | Vijayalakshmi *et al.* (2010), Naruoka *et al.* (2012) |
|  | Canopy persistence |  |  |  | Time to plant maturity |  |  |  | Vis. inspection | Adu *et al.* (2011) |
| Total Greenness | GA<sub>accum</sub>, Chl<sub>accum</sub>, LAUC, Sgint, TotalAUC |  |  |  | Greenness from anthesis to senescence completion Area under curve |  |  |  | Graphical calculation | Kumar *et al.* (2010), Lopes and Reynolds (2012), Christopher *et al.* (2014, 2016, 2018), Pinto *et al.* (2016), Xie *et al.* (2016) |
|  | Mean |  |  |  | Average flag leaf score over scoring duration |  |  |  | Mean | Adu *et al.* (2011) |
|  | p0, N<sub>green_max</sub>, MaxChl, MaxNDVI |  |  |  | Minimum senescence score or max greenness |  |  |  |  | Bogard *et al.* (2011), Christopher *et al.* (2014, 2018), Xie, *et al.* (2016), Kitonyo *et al.* (2017), Alhabbar *et al.* (2018) |
| Dynamic Measures | 50%G, TFN50*, MidS, 50 % senescence |  |  |  | Time to loss 50 % maximum greenness |  |  |  |  | Vijayalakshmi *et al.* (2010), Christopher *et al.* (2014, 2016, 2018), Kitonyo *et al.* (2017) |
|  | SG100 - SG1100 |  |  |  | Estimated NDVI from 100 to 1100 d °C post anthesis as 100 d °C intervals |  |  |  |  | Christopher *et al.* (2014, 2016) |
|  | %GFLA |  |  |  | Change in percentage green flag leaf area between anthesis and scoring dates |  |  |  | Raw data | Verma *et al.* (2004) |
|  | 75%G |  |  |  | Time to 25 % senesced |  |  |  |  | Vijayalakshmi *et al.* (2010) |
|  | 25%G, Days to Flag Leaf Senescence |  |  |  | Time to 75 % senesced |  |  |  |  | Vijayalakshmi *et al.* (2010), Naruoka *et al.* (2012) |

CIMMYT crop ontology defines the ‘Staygreen’ trait (CO_321:0000059) as the ‘Ability of the plant to remain/maintain green leaves, stems and spikes at the time of senescing’ (Shrestha *et al*., 2012). However, only functional staygreen phenotypes are considered useful due to their association with prolonged or enhanced photosynthetic activity, compared to cosmetic types in which chlorophyll catabolism is impaired (Gregersen *et al*., 2013; Thomas and Ougham, 2014). Unfortunately, senescence phenotyping efforts often concentrate on recording changes in leaf, or canopy, colour and not the accompanying developmental and physiological changes, potentially favouring identification of cosmetic staygreens, opposed to useful types.

A direct correlation between green canopy and grain fill duration is frequently assumed, which experiments by Wiegand and Cuellar (1981) and Gelang *et al*. (2000) confirm. However, when studying senescence this relationship is rarely explicitly tested, although Pinto *et al*. (2016) and De Souza Luche *et al*. (2017) report a positive correlation between flag leaf and grain fill duration, r = 0.16 to 0.7, *P* < 0.01. Loss of glume colour and peduncle ripening are also associated with changes in grain development, coinciding with GS87, the timepoint at which dry grain weight is maximal (Pask *et al*., 2012). Ear photosynthesis contributes to between 40 to 80 % of grain carbohydrate (Zhou *et al*., 2016), with ears supplying 1.87 × more nitrogen compared to flag leaves (Barraclough *et al*., 2014). Additionally, peduncle senescence has important implications regarding the delivery of flag-leaf derived photosynthates. Peduncles act as conduits and stores for transient starch and sugars, facilitating their remobilisation into the grain, whilst carbohydrates within peduncle tissue help maintain hydraulic conductance (Raven and Griffiths, 2015). If peduncles senesce in advance of leaves then the photosynthates associated are unable to reach the grain. Together, this illustrates the need for ear and peduncle phenotyping, alongside recording of grain filling dynamics when studying senescence, with Table 2 listing methods adopted.

**Table 2.**
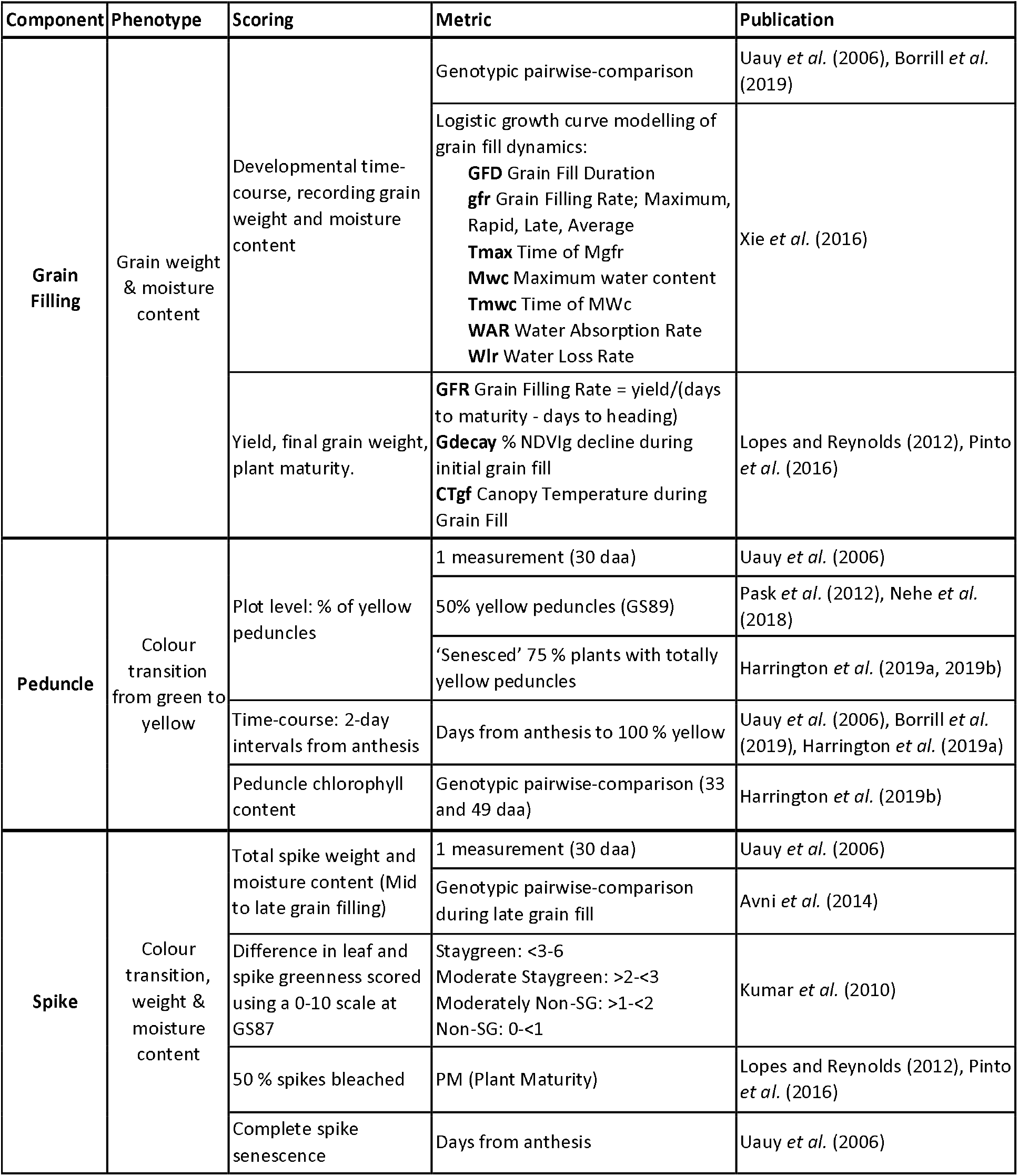
Senescence scoring efforts should not be limited to leaves. Senescence is a whole plant process involving remobilisation of resources into grain, for which studying multiple organs have been used to aid our understanding.

| Component | Phenotype | Scoring | Metric | Publication |
| --- | --- | --- | --- | --- |
| Grain Filling | Grain weight & moisture content | Developmental time-course, recording grain weight and moisture content | Genotypic pairwise-comparison | Uauy *et al.* (2006), Borrill *et al.* (2019) |
|  |  |  | Logistic growth curve modelling of grain fill dynamics:<br>**GFD** Grain Fill Duration<br>**gfr** Grain Filling Rate; Maximum, Rapid, Late, Average<br>**Tmax** Time of Mgfr<br>**Mwc** Maximum water content<br>**Tmwc** Time of MWc<br>**WAR** Water Absorption Rate<br>**Wlr** Water Loss Rate | Xie *et al.* (2016) |
|  |  | Yield, final grain weight, plant maturity. | **GFR** Grain Filling Rate = yield/(days to maturity - days to heading)<br>**Gdecay** % NDVIg decline during initial grain fill<br>**CTgf** Canopy Temperature during Grain Fill | Lopes and Reynolds (2012), Pinto *et al.* (2016) |
| Peduncle | Colour transition from green to yellow | Plot level: % of yellow peduncles | 1 measurement (30 daa) | Uauy *et al.* (2006) |
|  |  |  | 50% yellow peduncles (GS89) | Pask *et al.* (2012), Nehe *et al.* (2018) |
|  |  |  | 'Senesced' 75 % plants with totally yellow peduncles | Harrington *et al.* (2019a, 2019b) |
|  |  | Time-course: 2-day intervals from anthesis | Days from anthesis to 100 % yellow | Uauy *et al.* (2006), Borrill *et al.* (2019), Harrington *et al.* (2019a) |
|  |  | Peduncle chlorophyll content | Genotypic pairwise-comparison (33 and 49 daa) | Harrington *et al.* (2019b) |
| Spike | Colour transition, weight & moisture content | Total spike weight and moisture content (Mid to late grain filling) | 1 measurement (30 daa) | Uauy *et al.* (2006) |
|  |  |  | Genotypic pairwise-comparison during late grain fill | Avni *et al.* (2014) |
|  |  | Difference in leaf and spike greenness scored using a 0-10 scale at GS87 | Staygreen: <3-6<br>Moderate Staygreen: >2-<3<br>Moderately Non-SG: >1-<2<br>Non-SG: 0-<1 | Kumar *et al.* (2010) |
|  |  | 50 % spikes bleached | PM (Plant Maturity) | Lopes and Reynolds (2012), Pinto *et al.* (2016) |
|  |  | Complete spike senescence | Days from anthesis | Uauy *et al.* (2006) |

Recently, staygreen traits have received renewed interest due to their potential ability to increase yield and stress tolerance (Gregersen *et al*., 2013; Jagadish *et al*., 2015). For example, multiple studies report the indirect selection of staygreen traits over the previous 50 years has helped sustain grain number improvement (Adu *et al*., 2011; Kitonyo *et al*., 2017; Voss-Fels *et al*., 2019). Modelling of wheat ideotypes using 2050 climate change predictions weights staygreen traits highly, estimating associated yield benefits of 28-37 % and 10-23 % for Spanish, and Central and Eastern European growing regions respectively (Senapati *et al*., 2019). Under stress, Chapman *et al*. (2020) reported a positive relationship between delayed senescence and grain weight improvement of *NAM-1* EMS mutants, which could relate to elevated ABA levels enhancing carbon remobilisation (Distelfeld, Avni and Fischer, 2014). However, not all staygreen phenotypes are the same, with senescence dynamics a product of differences in the onset, rate, duration or initial chlorophyll content (Figure 2). Xie *et al*. (2016) hypothesise a delay in onset, coupled with a rapid rate, of senescence maximises remobilisation efficiency, reporting rapid grain fill rate and TGW as correlated, r = 0.63 to 0.77, *P* < 0.01. Conversely, Gelang *et al*. (2000) report senescence duration as the greatest contributing factor to grain fill, with traits highly associated, R^2^ = 0.989. In combination, elucidating the optimal combination of senescence dynamics under differing conditions is required to identify their associated target breeding environments and stimulate staygreen trait adoption.

**Figure 2.**
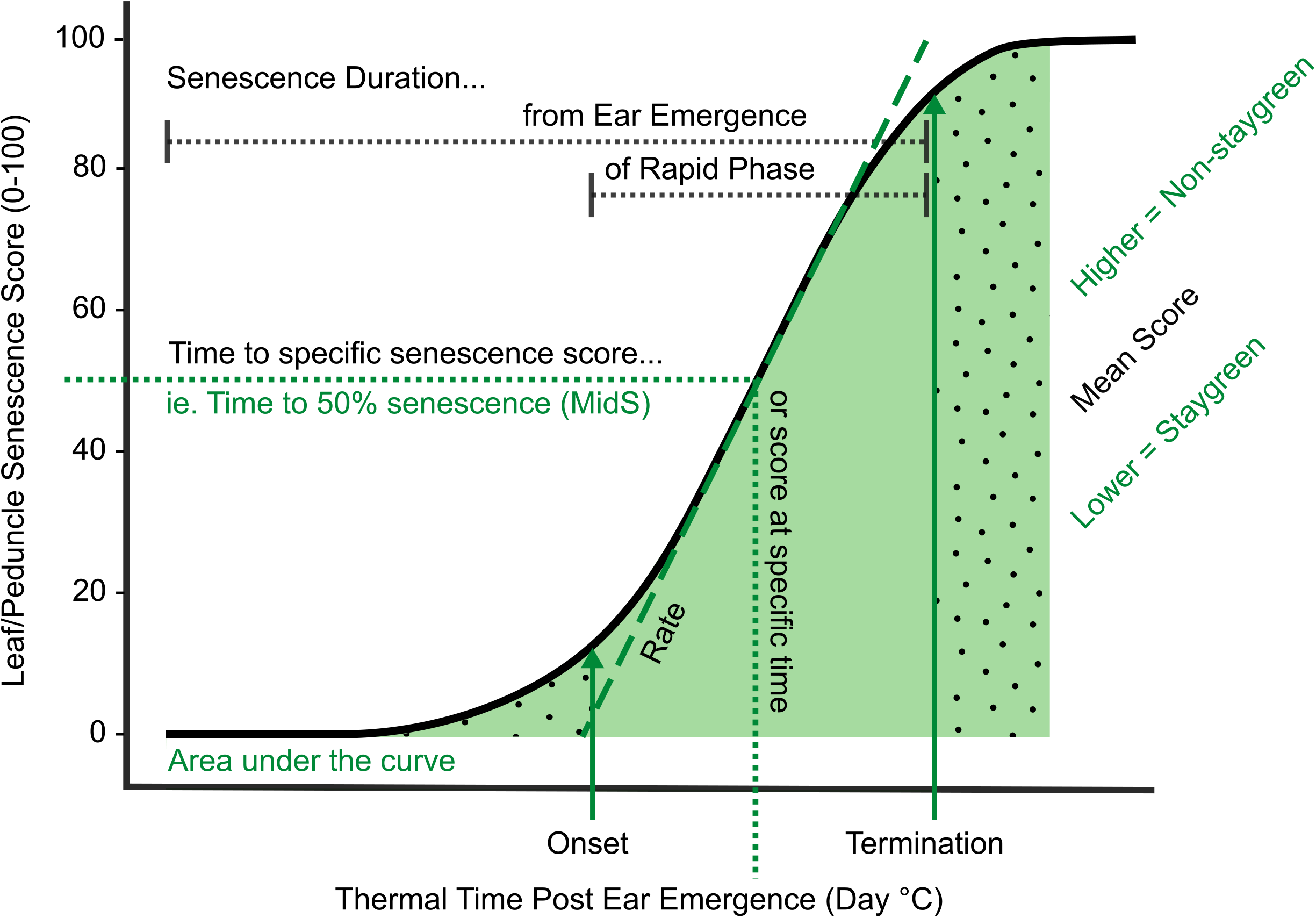
Key parameters used in quantification and characterisation of senescence variation. Metrics relating to onset, rate and duration of senescence, alongside total greenness (area under the curve) and dynamic measures (including MidS) can be calculated from time-course data (Table 2). Onset of senescence marks the transition between the initial lag phase (dotted fill, right) and rapid phase of senescence (solid fill, centre). As senescence nears completion the senescence rate decreases, resulting in a final lag phase (dotted fill, left). To compare senescence progression of different lines time to different senescence scores (MidS), an overall mean or total greenness can be calculated.

For studies investigating a limited number of lines, plotting and visual comparison of senescence time-course data may be sufficient to identify and characterise senescence variation. However, when assessing senescence of segregating RIL populations, or a diversity set, visual discrimination of individual senescence profiles is challenging (Figure 1). Transformation of senescence curves into a series of well-defined metrics aids characterisation of individual lines, allows senescence dynamics to be described, and permits performance of quantitative analysis, including QTL mapping. Unfortunately, multiple studies apply their own methods to derive senescence metrics, leading to varying definitions of apparently similar terms (Table 1). In absence of consistent senescence phenotyping approaches results from different studies cannot be directly compared, preventing interpretation of significant genotypic and environmental variation (Verma *et al*., 2004; Pask *et al*., 2012; Christopher *et al*., 2014).

Recognising the disparity in senescence phenotyping methods present in the literature, we reviewed, used, and developed a range of methods that successfully capture senescence variation observed amongst two segregating RIL populations. Although duration and onset of senescence are the metrics most used to describe senescence dynamics (Table 1), these can fail to capture process dynamism and source to sink relationships. Simultaneously scoring leaf and peduncle senescence, alongside monitoring changes in grain development, improved our understanding of senescence processes at a whole plant level. We also identified the need for singular metrics capable of discriminating between staygreen and non-staygreen types, preferably in absence of time-course phenotyping, to increase efficiency of in-field phenotyping and selection. Here, we compile resources we referred to when scoring senescence under field conditions, providing our own insights following successful mapping of *NAM-1* homoeologues, which are known senescence regulators (Chapman *et al*., 2020). Here we aim to equip researchers and breeders alike with the appropriate knowledge to guide future phenotyping strategies, support trait genetic mapping and selection, and help to inform weighting of senescence traits in breeding, genomic selection or other applications.

## Materials and Methods

### Plant Material

Phenotypic data relates to two *Triticum aestivum* cv. Paragon EMS staygreen mutants and associated Recombinant Inbred Line (RIL) populations. ‘Staygreen A’ and ‘Staygreen B’ refers to mutant lines 1189a and 2316b, identified as encoding missense mutations in known senescence regulators *NAM-A1* (T159I) and *NAM-D1* (G151), respectively (Chapman *et al*., 2020). ‘Non-staygreen’ refers to the parental *Triticum aestivum* cv. Paragon. To develop segregating RIL populations mutants were crossed to cv. Paragon and F4 populations developed through SSD, n ≥ 85.

### Field Experiments

Phenotyping of F4 RIL populations was conducted under field conditions between 2016 and 2018. Experiments were performed at Church Farm, Bawburgh (52°38’N 1°10’E), JIC, as described previously (Chapman *et al*., 2020). In brief, 36 to ≥ 75 RILs were sown per population per year as unreplicated 1 m^2^ plots (2016) or replicated 6 m^2^ plots (n ≥ 2; 2017 & 2018). Seeds were sown on 26/10/2016, 26/10/2017 and 12/10/2018 at a rate of 250 to 300 seeds m^−2^. Replicated experiments followed a randomised complete block design, with control plots of cv. Paragon, Soissons, 1189a and 2316b mutant lines randomly sown throughout. Seed used for experiments was produced during the multiplication of RIL populations in 2015 or resulted from the previous year. Soil at Church Farm is described as sandy loam overlying alluvial clay. Supplemental irrigation was applied in 2017, otherwise trials were naturally rainfed. Fertiliser was applied over three occasions from late February to the end of April, totalling 214 kg N ha^−1^ and 62 kg SO_3_ ha^−1^ in 2017, and 228.5 kg N ha^−1^ and 62 kg SO_3_ ha^1^ in 2018. Plots received standard fungicide and herbicide treatment.

### Phenotypic assessment

Ear emergence (GS55) was recorded as when 50 % of ears emerged halfway from the flag leaf across the plot (Zadoks *et al*., 1974). Leaf and peduncle senescence were visually scored from ear emergence to maturity every 2-3 days using a 0-100 scale (intervals of 5) (Figure 3).

**Figure 3.**
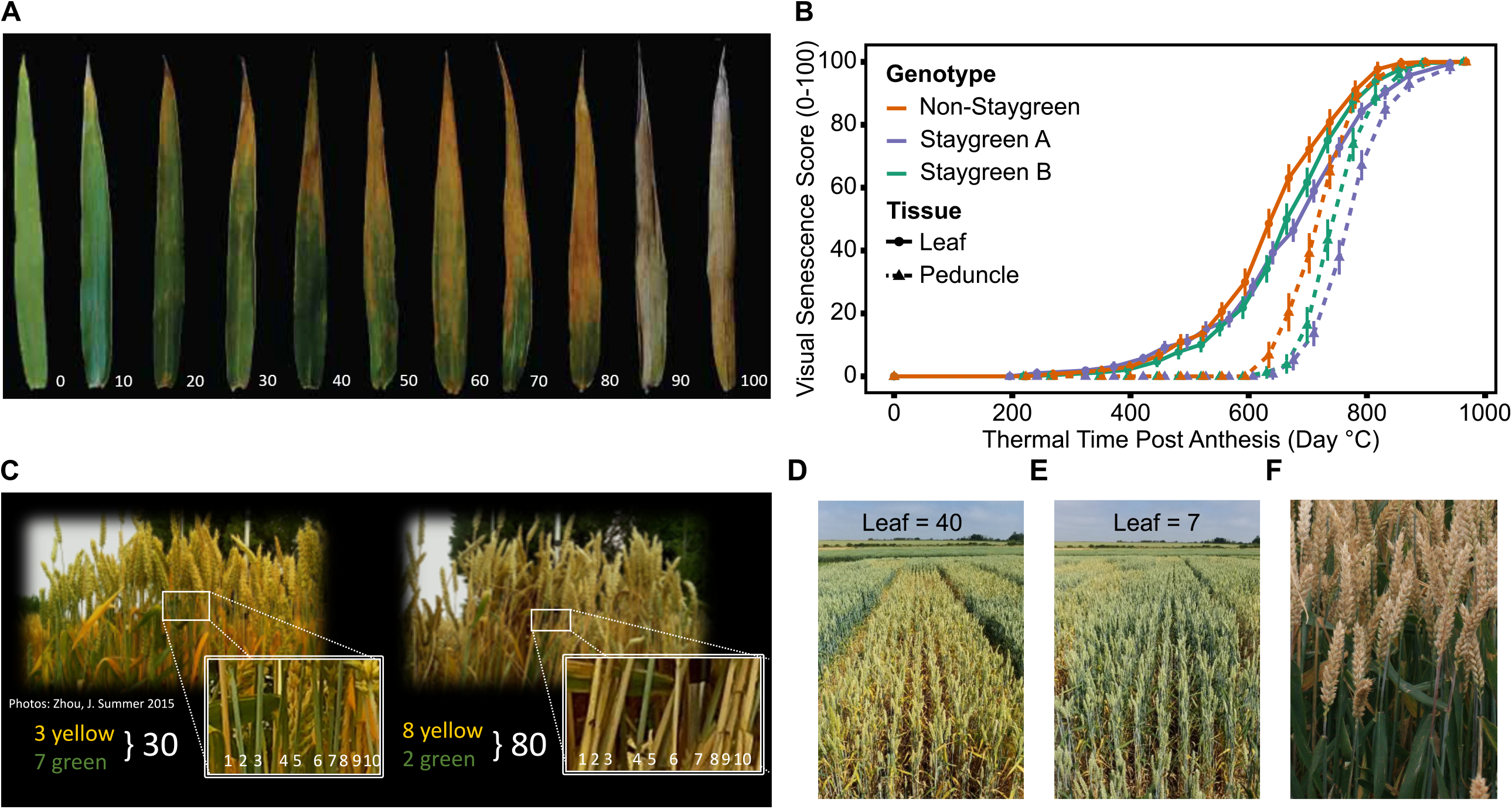
Simultaneous scoring of flag leaf, peduncle and ear senescence allows the whole plant nature of senescence to be considered. Compared to flag leaf senescence **(A)**, scoring of peduncle senescence **(C)** is less subjective, and respective senescence profiles were found to reinforce one another **(B)**. Senescence of multiple plants was visually assessed and scored using a 0-100 scale **(A** & **C)** at the plot level **(D & E)**, taking care to avoid edge effects or diseased plants. When observed, asynchronous senescence phenotypes **(F)** were recorded and may indicate impaired remobilisation efficiency.

Flag leaf senescence was scored as the proportion of flag leaf yellowing; a score of 5 indicating leaf tip necrosis, and 100 complete senescence (Figure 3A). To avoid edge effects, flag leaves of multiple plants within plot centres were assessed together to derive an overall plot score (Figure 3D-E). Instances of plot heterogeneity resulting from disease, soil gradients or damage were also recorded and subsequently referred to for the purposes of outlier detection. To reduce systematic error plots were scored in the same orientation and direction on each visit, with scoring in direct sunlight avoided due to increased difficulty of identifying plot differences.

Peduncle senescence was scored as the percentage of plants for which the top 5 cm of peduncle tissue had transitioned from green to yellow. Compared to leaves, peduncles senesce evenly along their length, and the phase of rapid senescence is shorter (Figure 3B). This rapid colour transition enables peduncles to be scored as either completely green or completely yellow, increasing objectivity of assessment. To provide a plot score 3-4 batches of 10 tillers were assessed and percentage yellow derived (Figure 3C).

Previously, grain filling experiments were performed for Staygreen A, Staygreen B and Non-staygreen parental cv. Paragon, in which a significant grain fill extension was reported for both staygreen lines (Chapman *et al*., 2020). To identify any potential association between grain filling and senescence phenotypes, grain maturity of RIL populations was scored according to the Zadoks scale with reference to the ‘Wheat Growth Guide’ (Zadoks *et al*., 1974; AHDB, 2018). 2-3 immature grains, from two plants per plot, were subject to thumbnail impressions or squashed between finger and thumb to determine developmental stage, with observations recorded using a 1-4 scale (hard to soft) or Zadoks growth stages, GS79-GS93 (Milky dough to ripe, grain loosening in the daytime) (AHDB, 2018). When observed, differences in leaf and ear senescence were recorded, which in extreme cases manifested themselves as ‘green leaf, ripe ear’ phenotypes (Figure 3F). Through recording leaf, peduncle, grain and ear phenotypes one can understand the whole plant nature of senescence, providing insights into resource remobilisation and source to sink relationships. Table 2 lists senescence phenotyping approaches applied by other studies.

### Derivation of Senescence Metrics

To quantify and interpret senescence dynamics multiple senescence metrics can be derived (Figure 2). Leaf and peduncle senescence scores were plotted against thermal time post ear emergence GS55 (°C day) to standardise for heading variation and associated differences in temperatures exposed. Mean daily temperature was calculated using daily minimum and maximum temperatures and summated over time in days, with GS55 corresponding to 0 °C days. Metrics used to describe senescence patterns fall into five categories, corresponding to onset, rate, duration, total greenness, or a dynamic measurement of senescence, however definitions of these terms alongside their method of calculation vary (Table 1).

The sigmoidal progression of senescence facilitates curve modelling, whereupon senescence is divided into three phases: an initial lag phase, a rapidly senescing phase, and a final lag maturation phase (Table 1). To calculate rate, onset and duration of senescence we initially applied the model described by Bogard *et al*. (2011) and Gaju *et al*. (2011) to our data. A comparison of raw and modelled curved found calculated metrics were of limited use identifying a tendency of the model to overfit the data, whilst infrequent scoring led to inaccurate calculation of inflection points.

Due to problems encountered when curve modelling, we reviewed the definitions of commonly calculated metrics (Table 1) to inform derivation of senescence metrics from raw data (Table 3). Concordant with Christopher *et al*. (2014, 2016, 2018), Xie *et al*. (2016), Alhabbar *et al*. (2018) and Kitonyo *et al*. (2017) we define onset of senescence as the ‘start of rapid senescence phase (Leaf score ∼10-15)’ (Table 3). Similar to Christopher *et al*. (2014, 2016), Xie *et al*. (2016) and Nehe *et al*., 2018), we define termination of senescence as the ‘time at which maximum senescence score (>90) is first recorded’ (Table 3). Senescence duration was calculated from both ear emergence and onset of senescence (Figure 2), with the latter providing an indirect measure of senescence rate (Table 3). To capture senescence dynamism we derived times to different flag leaf senescence scores (Table 3), similar to metrics MidS (midpoint of senescence) and 75%G utilised by Vijayalakshmi *et al*. (2010) and others (Table 1). Assuming senescence progresses linearly, time points corresponding to scores above and below the ‘target’ senescence score, ie 70, are identified with time divided proportionately to estimate time elapsed (Figure 4). Calculating the mean senescence score over the scoring period provides an assessment of overall greenness (Nehe *et al*., 2018), and was calculated separately for both flag leaf and peduncle senescence (Table 3).

**Table 3.** Senescence metrics derived for quantification and qualification of time-course senescence profiles. Shading indicates tissue phenotyped.

| Metric | Abbreviation | Tissue |  | Description | Interpretation |
| --- | --- | --- | --- | --- | --- |
|  |  | Flag Leaf | Peduncle |  |  |
| Mean Senescence | **MeanLeaf**<br>**MeanPed** |  |  | Mean senescence score for total scoring period | Lower = Staygreen,<br>Higher = Non-staygreen |
| Onset of Senescence | **Onset** |  |  | Start of rapid senescence phase (Leaf score ~10-15) | Delay in senescence |
| Duration of rapidly senescing phase | **LeafSenDur**<br>**PedSenDur** |  |  | Time to maximum senescence score (Leaf > 90) – onset (Leaf ~10-15) | Senescence rate indicator<br>Lower = Faster |
| Ear Emergence to terminal senescence | **EEtoLeafSen**<br>**EEtoPedSen** |  |  | Time at which maximum senescence score (>90) first recorded | Senescence duration;<br>Longer = Staygreen,<br>Shorter = Non-staygreen |
| Thermal time to leaf senescence score 25, 30, 40..., n | **TT25, TT30,**<br>**TT40..., TT80** |  |  | Time to different flag leaf senescence scores, as interpolated from time-course data (Figure 4) | Dynamic measure;<br>Later = Staygreen,<br>Earlier = Non-Staygreen |

**Figure 4.**
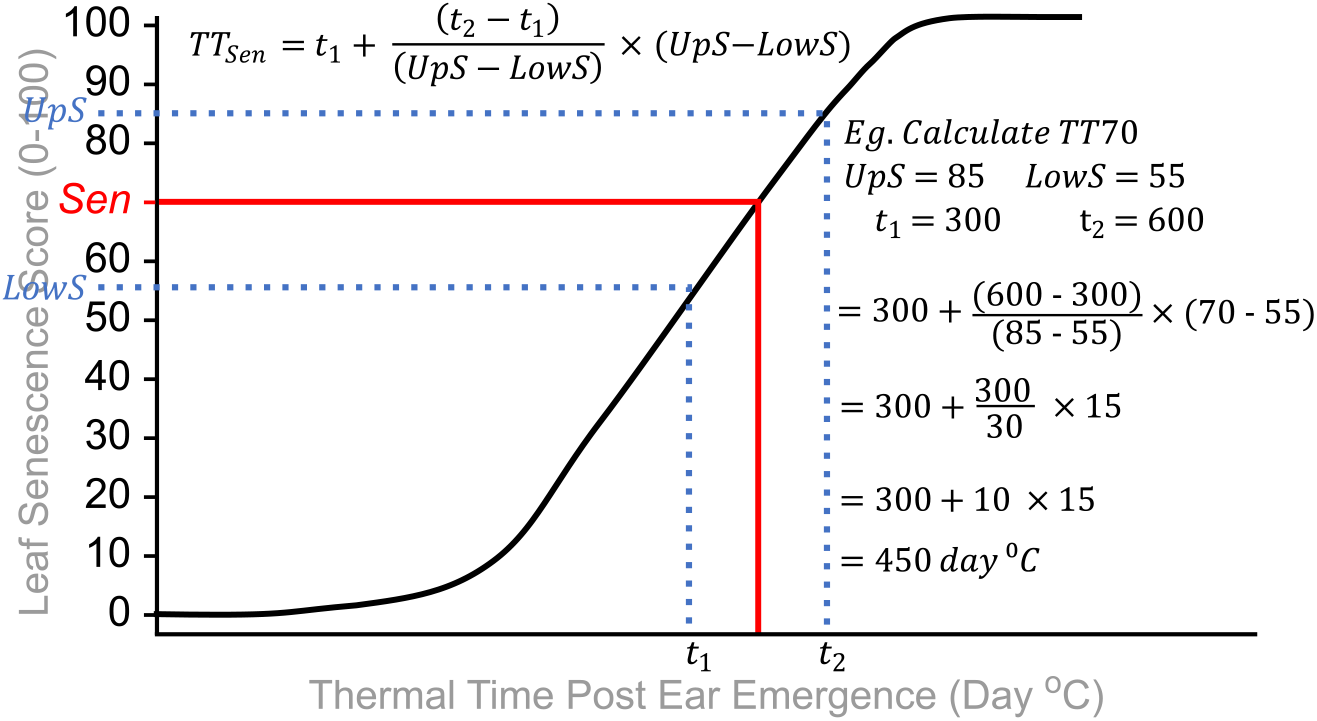
Calculation of TT70, a dynamic senescence metric. Senescence scoring dates are converted into thermal time from ear emergence (Day °C) to standardise for heading-date variation. A ‘target’ senescence score is specified *(Sen)*, and time points for scores below (t_1_, *LowS)* and above (t_2_, *UpS)* this identified. Assuming senescence progresses linearly, time elapsed between senescence scores is divided proportionately to estimate the time to the target score (*TT*_*Sen*_).

### Data Analysis

Data analysis was performed using R (version 3.5.2) (R Core Team, 2018) within RStudio (RStudio team, 2015), and data manipulated using packages ‘data.table’ (Dowle *et al*., 2019), ‘dplyr’ (Wickham *et al*., 2018), ‘plyr’ (Wickham, 2015) and ‘tidyr’ (Wickham *et al*., 2019). Senescence metrics were derived from raw senescence data in absence of spatial correction and means calculated per line when replicated. To assess heritability and accuracy of senescence scoring Spearman’s rank correlations were calculated and results visualised using ‘ggpubr’ (Kassambara, 2018). To illustrate the discriminatory power of senescence metrics phenotype × genotype plots were constructed using package ‘r/qtl’ (Arends *et al*., 2010), with other graphs produced using package ‘ggplot2’ (Wickham *et al*., 2018). To determine significance of phenotypic differences linear mixed modelling was performed using packages ‘lme4’ (Bates *et al*., 2019) and ‘lmerTest’ (Kuznetsova *et al*., 2017), with replicate, row, column and *NAM-1* genotype treated as fixed effects, and RILs per population as random. Tukey-Post hoc tests were performed using package ‘lsmeans’ (Lenth, 2018).

## Results

### The Parallel Progression of Peduncle Senescence

Few publications report the use of peduncle senescence phenotyping when studying senescence regulation (Table 2). In 2017 and 2018 we conducted time-course senescence scoring of leaf and peduncle tissue (Figure 3). We identified peduncle senescence is initiated after flag leaf senescence, with senescence profiles found to reinforce one another, aiding differentiation of lines (Figure 3B; Chapman *et al*. (2020)). Compared to flag leaf senescence, peduncle senescence occurs over a shorter time-period and individual scores demonstrate less variation, indicated by their comparatively smaller error bars (Figure 3B).

### Methods for Accurate Capture of Senescence Variation

Between 2016 and 2018 RILs segregating for two independent senescence traits were phenotyped under field conditions. Plotting senescence data against thermal time enabled calculation of senescence metrics, illustrating the effect of environment on senescence regulation. For example, between 2016 to 2018 Ear Emergence to Terminal Flag Leaf Senescence (EEtoLeafSen) scores ranged from 825 [792, 858] to 1156 [1090, 1220] °C d for the ‘Non-staygreen’ parent, a difference of 0020331 °C d.

To determine the stability of senescence phenotypes, and accuracy of phenotyping methods, year-pairwise Spearman’s rank correlations were calculated for a range of senescence metrics. We found the magnitude and significance of between-year phenotypic correlations relate to penetrance and stability of parental senescence phenotypes. For example, Staygreen A RILs segregate for a relatively extreme staygreen phenotype (Figure 3B) and year-pairwise phenotypic correlations ranged from *r* = 0.39 to 0.91, *P* ≤ 0.02 (Supplemental Table 1). Conversely, Staygreen B RILs segregate for a milder staygreen phenotype and year-pairwise phenotypic correlations were lower, ranging from *r* = 0.3 to 0.68, *P* ≤ 0.03 (Supplemental Table 1). However, correlations were significant for all three year-pairs for between 2 to 9 metrics, *P* ≤ 0.03, indicating trait heritability, environmental stability, and accurate phenotyping of RILs (Supplemental Table 1). Insights from such analysis can direct in-field phenotyping approaches, informing which senescence metrics to prioritise when conducting phenotypic selection and forward genetic screens.

As time-course phenotyping is time consuming and laborious, we identified the need for a single senescence metric capable of discriminating senescence types. Inspection of year-pairwise correlations identified metric TT70 as a potential candidate, with correlations of *r* = 0.78 to 0.84 and *r* = 0.37 to 0.59, *P* ≤ 0.02, reported for Staygreen A and Staygreen B RILs, respectively (Figure 5C, Supplemental Table 1). Alternatively, peduncle senescence derived metrics may better discriminate senescence phenotypes. Compared to metric TT70, year-pairwise correlations for MeanPed (Mean Peduncle Senescence) scores were greater, ranging from *r* = 0.62 (Staygreen B) to 0.91 (Staygreen A), *P* < 5 × 10^−6^ (Figure 5B), indicating greater environmental stability.

**Figure 5.**
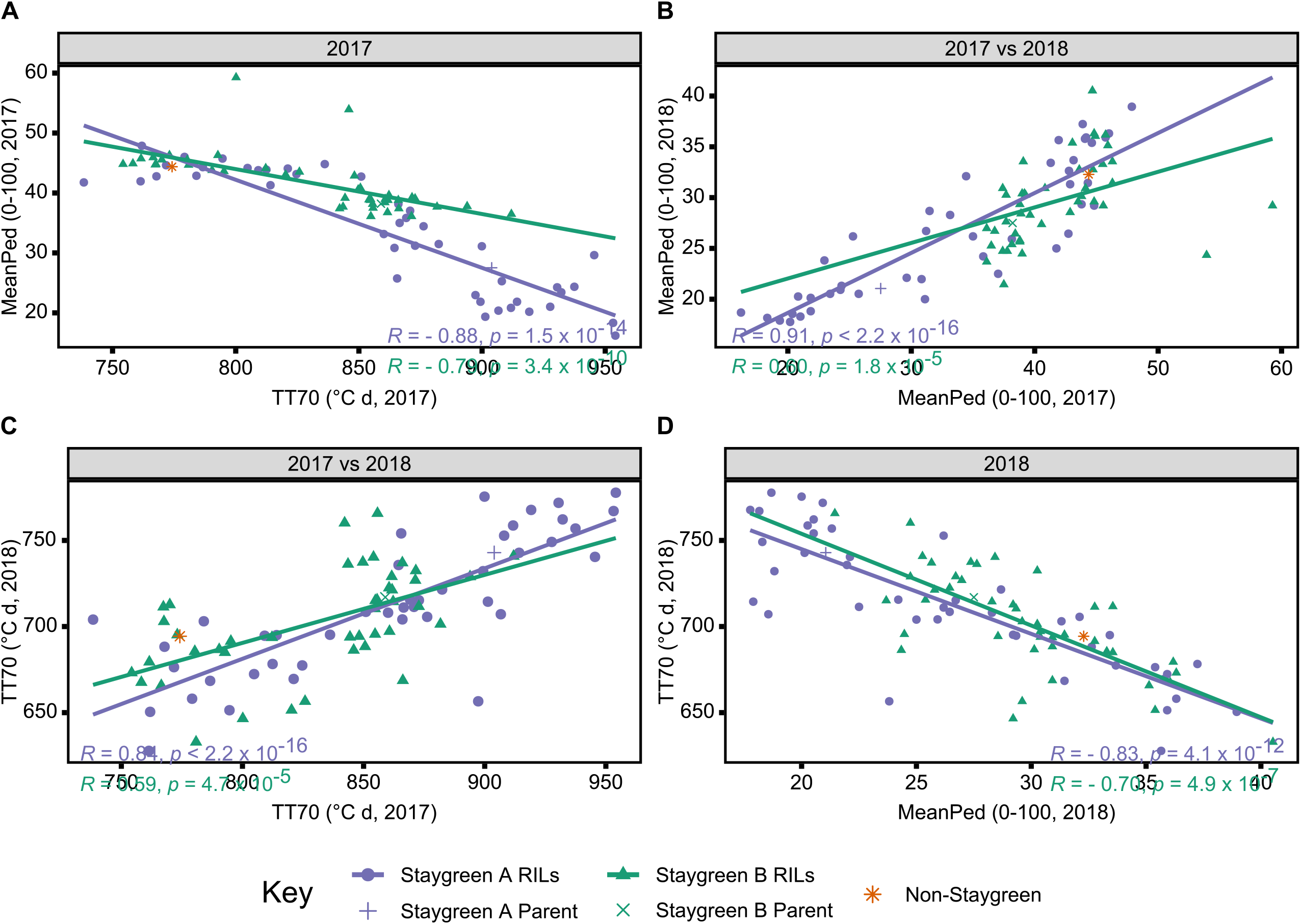
Environmental stability and concordance of metrics TT70 and MeanPed aids discrimination of staygreen and non-staygreen types. Between-year Spearman’s rank correlations for metrics MeanPed **(B)** and TT70 **(C)** range from *R* = 0.59 to 0.91, *P* < 0.0001, with metrics themselves negatively correlated, *R* = 0.7 to 0.88, *P* < 0.0001 **(A & D)**. Comparing TT70 and MeanPed values recording in 2017 **(A)** and 2018 **(B)** one can distinguish senescence variation, whereby ‘staygreen’ and ‘non-staygreen’ lines cluster in the bottom-right and top-left corners, respectively. Spearman’s rank correlations calculated for the 36-42 RILs per population grown at Church Farm, Norwich (JIG) between 2017 and 2018, plotting mean value per line, n ≥ 2. Staygreen A parent and RILs (purple), Staygreen B parent and RLs (green), recurrent parent (orange).

### Coupling MeanPed & TT70 Scores Aids Selection and Discrimination of Senescence Types

Calculation of year-pairwise phenotypic correlations illustrate our approach to senescence scoring and quantification is robust, but does it aid discrimination of senescence variation amongst lines? Correlation plots displaying mean TT70 or MeanPed scores recorded for individual RILs reveals their tendency to cluster into senescence types. For metric TT70, RILs clustered towards the bottom-left corner are considered ‘non-staygreen’, with those clustered towards the top-right ‘staygreen’ taking longer to senesce (Figure 5C). Conversely, lower MeanPed scores indicate the greater retention of green peduncle tissue, with RILs clustered towards the bottom-left considered ‘staygreen’ (Figure 5B).

The degree of separation between ‘non-staygreen’ and ‘staygreen’ clusters relates to the extremity of senescence phenotype. For example, the mean difference in TT70 scores between lines Staygreen A and B compared to the parental Non-staygreen line are 85.3 ± 34.4 °C d and 63.0 ± 27.8 °C d, respectively, with contrasting Staygreen A RILs clustering further apart (Figure 5C). However, greater robusticity of peduncle-derived senescence scores contributes to tighter clustering of RILs compared to TT70 (Figure 5B). In combination, metrics TT70 and MeanPed can be used to accurately discriminate senescence types, particularly in absence of multi-year phenotyping data as we found metrics to be highly correlated *r* = −0.7 to −0.83, *P* ≤ 4.9 × 10^−6^ (Figure 5A & D).

Proof of the utility of TT70 and MeanPed scores in discriminating senescence variation are results of mapping by bulk segregant analysis conducted for Staygreen A and Staygreen B (Chapman *et al*., 2020). Whilst different senescence metrics may better capture the range of senescence variation in different years, assessment of TT70 and MeanPed scores consistently identified the same RILs for which senescence was delayed, facilitating the selection of phenotypically contrasting bulks. Compared to metric MeanLeaf, the distribution of MeanPed scores recorded for staygreen A and B RILs was flatter, indicating peduncle derived senescence scores could better discern senescence variation (Figure 6).

**Figure 6.**
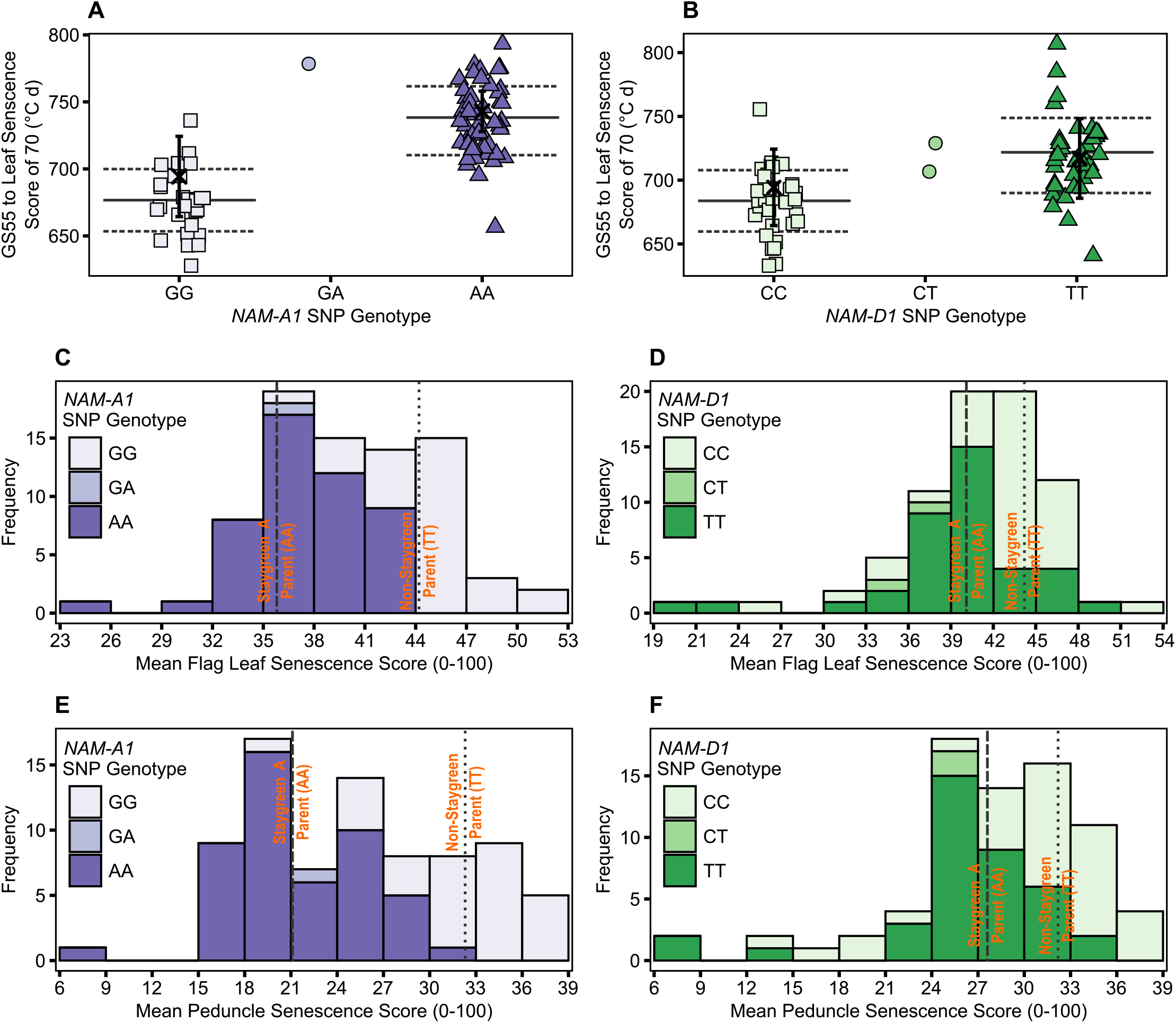
Illustrating the differential power of metrics TT70, MeanLeaf and MeanPed in discriminating senescence variation. Phenotypic differences were significant between homozygous RILs contrasting for mutations in NAM-A1 (Staygreen A) **(A, C, E)** and NAM-D1 (Staygreen B) **(B, D, F)** according to TT70 **(A & B)**, MeanLeaf (C **& D)** and MeanPed **(E & F)** scores, *P* ≤ 0.003. Metric MeanPed proved more informative when classifying senescence types compared to MeanLeaf due to the greater range and spread of scores. Phenotypic differences between parents were not significant, *P* > 0.05. **(A & B)** Scatterplot of TT70 scores against *NAM-1* genotype; genotypic group mean (solid line) ± SD (dotted line), Parental mean (black cross)± SD. **(C to F)** Bars display mean MeanLeaf and MeanPed score recorded for two RIL populations, n ≥ 75 (2 replicates) segregating for mutations in *NAM-A1* **(C, E)** or *NAM-D1* **(D, F);** Allelic combinations, GG/CC = homozygous non-staygreen, AA/ TT= homozygous staygreen A/staygreen B, GA/CT= heterozygous. Lines represent parental means (n > 5); Staygreen parents (dashed), Non-staygreen parent (dotted). Trial grown at Church Farm, Norwich (JIC), 2018.

However, although differences in leaf and peduncle senescence profiles of Staygreen A and B were typically significant relative to the non-staygreen line, *P* < 0.0001 to *P* = 0.11 (Chapman *et al*., 2020), differences in TT70 and MeanLeaf scores were not significant in 2018, *P* > 0.05, but were in 2016 and 2017, *P* < 0.05. Similarly, differences in MeanPed scores recorded for Staygreen A and the Non-staygreen parent were significant only in 2017, *P* < 0.01, whilst differences observed for Staygreen B were not, *P* > 0.05. Therefore, when differences in a senescence metric are not significant between lines under investigation one recommends their comparative assessment, just as we performed when classifying senescence variation amongst RILs during mapping of Staygreens A and B (Chapman *et al*., 2020). This approach worked successfully, as following trait mapping differences in TT70 and MeanPed scores recorded for homozygous RILs contrasting for mutations in *NAM-A1* or *NAM-D1* (Staygreen A and B, respectively) were significantly different in all three years, *P* < 0.05 (Figure 6).

### Determining the Association between Delayed Senescence and Grain Maturity

Previously, grain filling experiments conducted for Staygreens A & B found that grain moisture content remained elevated during the final 10-15 days of grain filling, *P* < 0.05 (Chapman *et al*., 2020). The number of time points for which grain moisture content of Staygreen A and Staygreen B were significantly greater compared to their common non-staygreen parent corresponded to the observed differences in onset of senescence, indicating a positive trait association (Chapman *et al*., 2020). Variation in grain development was detected amongst RIL subsets segregating for senescence traits grown in 2016 and 2017 (data not shown), however because Staygreens A & B were produced through EMS mutagenesis involvement of background mutations could not be discounted.

Conduction of grain filling experiments for entire RIL populations would have been unfeasible, however thumbnail impressions are routinely performed to assess grain development and maturity (Zadoks *et al*., 1974; AHDB, 2018). On 20^th^ July 2018 grain filling experiments found grain moisture content of Staygreens A & B was significantly elevated, *P* < 0.001 (Chapman *et al*., 2020). Thumbnail-based maturity scores ranged from GS87 to GS87-91 for the Staygreen A parent (n = 4), GS87-91 for the Staygreen B parent (n = 3), and GS92 for the Non-Staygreen parent (Figure 7). Grain maturity of some plots was recorded as being between two growth stages, GS87-91 and GS91-92, attributable to variation between main and secondary tillers.

**Figure 7.**
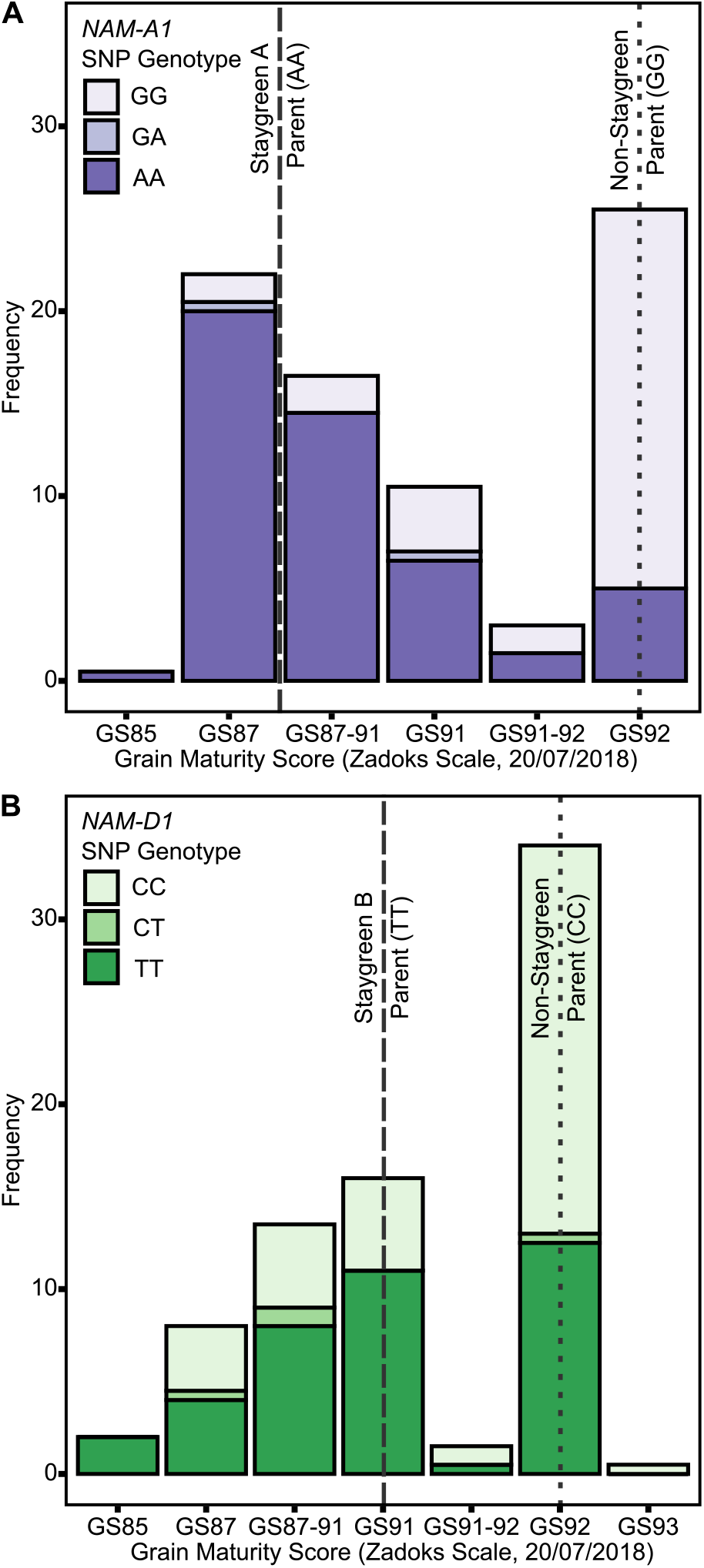
Staygreen phenotypes are positively associated with delayed grain maturation. Grain maturation of RIL populations segregating for senescence traits was assessed when significant differences in grain moisture content between parental lines were previously observed. Grain maturation of RILs homozygous for mutations in *NAM-A1* (Staygreen A) **(A)** and *NAM-D1* (Staygreen B) **(B)** was delayed, with scores starting from GS85 onwards, compared to the predominant score of GS92 for RILs homozygous for the non-staygreen allele. Grain scored according to the Zadoks scale (Zadoks et al., 1974), assisted by ‘The Wheat Growth Guide’ (AHDB, 2018). Bars chart RIL populations, n ≥ 75 (2 replicates) segregating for a mutation *NAM-A1* (Staygreen A) **(A)** or *NAM-D1* (Staygreen B) **(B);** Allelic combinations, GG/CC = homozygous non-staygreen, AA/TT= homozygous Staygreen A/Staygreen B, GA/CT= heterozygous. Lines represent parental means (n > 5); Staygreen parents (dashed), Non-staygreen parent (dotted).

Grain maturity scores recorded for RIL populations (n ≥ 75) ranged from GS85 (soft dough) to GS93 (Grain loosening in daytime) (AHDB, 2018), which when plotted against *NAM-1* SNP composition confirm senescence and grain filling traits are associated, Figure 7. On 20^th^ July 2018, a grain maturity score of GS92 was recorded for 71 % and 59 % of RILs homozygous for the ‘Non-staygreen’ *NAM-A1* (GG) or *NAM-D1* (CC) allele, respectively (Figure 7). In contrast, 73 % of RILs homozygous for the Staygreen A *NAM-A1* allele (AA) recorded a grain maturity score of GS87-91 or below (Figure 7A). Differences in grain maturity were subtler for Staygreen B, and scores of GS85 to GS87-91 and GS91 were recorded for 37 % and 29 % of RILs homozygous for the *NAM-D1* allele (TT), respectively (Figure 7B). Previous grain filling experiments identified the relative timing of differences in grain development for Staygreens A & B (Chapman *et al*., 2020), and based on these results thumbnail impression-based scoring can quickly and accurately capture such differences. Assessing visual leaf and peduncle senescence together with grain maturation could provide a method to separate cosmetic and functional staygreen phenotypes. Furthermore, through recording visual leaf and peduncle senescence scores when grain maturity is reached one can identify if resources are being efficiently remobilised into the grain (Figure 3F).

## Discussion

### Perks of Peduncle Senescence Scoring

When mapping the *GPC-B1* locus, Uauy *et al*. (2006a) reported changes in peduncle colour as being linked. This resulted in the identification of the NAC transcription factor and senescence regulator *NAM-B1* (Uauy *et al*., 2006b). Subsequent reverse-genetic studies investigating the role of *NAM-1* homoeologues and paralogs in senescence regulation independently confirm the utility of peduncle phenotyping, with peduncle phenotypes accurately capturing senescence variation (Cantu *et al*., 2011; Avni *et al*., 2014; Pearce *et al*., 2014; Borrill *et al*., 2019; Harrington *et al*., 2019a). Compared to flag leaf senescence, peduncle senescence is initiated later and progresses rapidly, with visual yellowing of peduncles first observed when flag leaf senescence scores approach ∼50 (Figure 3B). Unlike leaves, peduncles senesce evenly along their length making plot-level assessment objective (Figure 3), with scores typically only confounded by Barley Yellow Dwarf Virus-associated anthocyanin production (Livingston *et al*., 1998).

In agreement with Borrill *et al*. (2019) and Harrington *et al*. (2019a, 2019b) we report greater environmental stability of peduncle, as opposed to flag leaf, senescence phenotypes (Figure 5B; Figure 5C). For example, when characterising *Triticum turgidum* cv. Kronos *NAM-A1* mutants, Harrington *et al*. (2019a) reported peduncle senescence as consistently delayed under both field and glasshouse conditions, *P* < 0.05, whereas flag leaf senescence was not, *P* > 0.05 (glasshouse), *P* < 0.001 to *P* > 0.05 (field). Regarding our own lines, we found peduncle and flag leaf senescence derived metrics were highly correlated, *R* > 0.7, *P* < 0.001 (Figure 5), illustrating both phenotyping approaches accurately capture senescence variation. Together, increasing our reliance on peduncle senescence phenotyping may reduce the need for prolonged time-course assessment, but to prevent this short window from being missed flag leaf senescence requires monitoring.

### Quantitative to Qualitative: Selecting Senescence Types

To quantify senescence, we initially calculated the mean flag leaf senescence score for the total scoring period (MeanLeaf). Comparing MeanLeaf scores of individual lines against their flag leaf senescence profiles revealed the metric poorly captured senescence dynamism. Similarly, when evaluating 14 Australian wheat cultivars Kitonyo *et al*. (2017) reported cv. Heron as early but slow to senesce, and cv. Justica CL Plus as greener overall and rapidly senescing, however mean senescence scores may be similar (Figure 8). Using an alternative approach, Kitonyo *et al*. (2017) applied a logistic regression to time-course NDVI measurements to quantify senescence, finding maximum NDVI scores (near flowering) increased with year of cultivar release. In wheat, use of the metric ‘mean senescence’ is rare (Table 1), suggesting its limited utility, whereas in maize Ziyomo *et al*. (2013) and Parajuli *et al*. (2018) successfully used mean scores to characterise stress responses of lines grown under different agronomic and cropping systems. Therefore, when quantifying senescence thought should be given to the specific pattern, or phase of senescence depicted, which may vary between systems.

**Figure 8.**
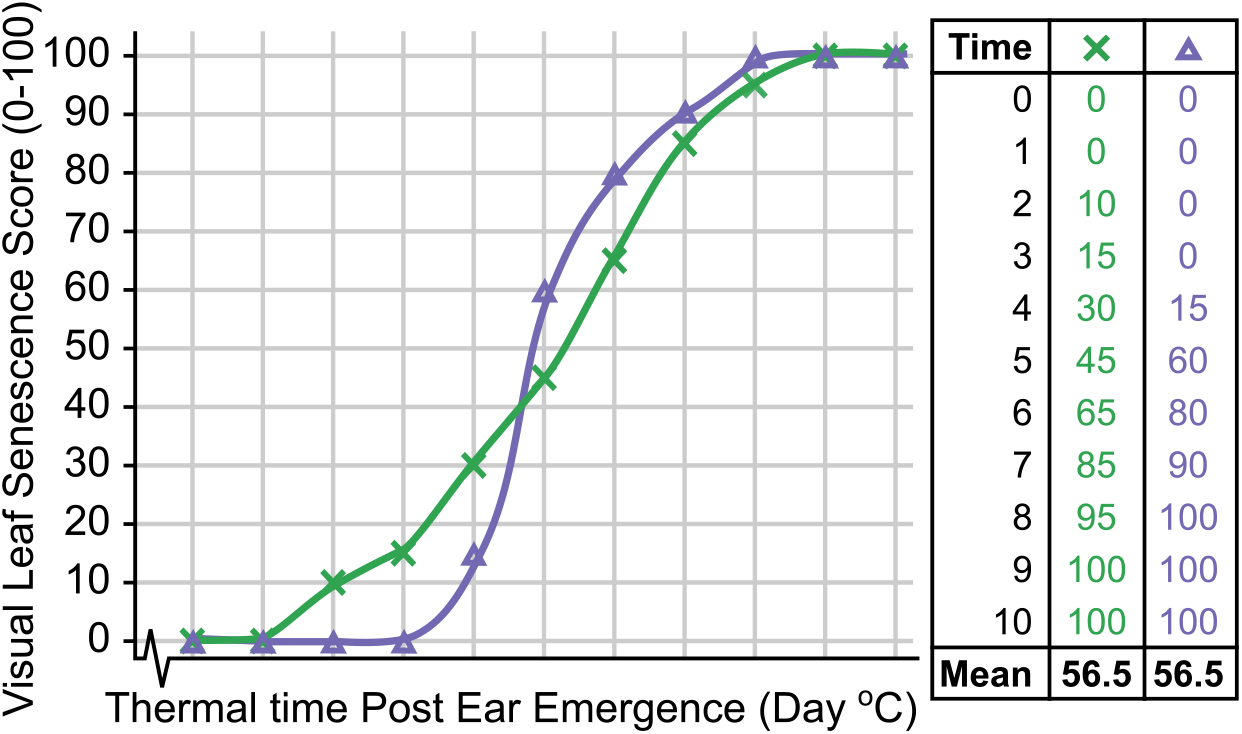
Mean Senescence Scores cannot always distinguish senescence variation. Senescence duration of line × (green) is prolonged, whilst onset of senescence is delayed and senescence rate rapid for line Δ (purple), but their mean senescence scores are the same.

In our study the metric Thermal Time to Flag Leaf Senescence Score of 70 (TT70) proved most informative when discriminating senescence types. Unlike calculation of mean leaf senescence scores, estimation of the thermal time taken to reach different flag leaf senescence scores (TT25, TT30…) was less affected by pre-existing leaf damage resulting from nitrogen splash, leaf tip necrosis or other damage. Pre-existing leaf damage appears as ‘noise’ during the early stages of senescence, however following the onset of senescence clear differences between lines emerge when flag leaf senescence scores range from 40 to 70 (Figure 1B). Depending on extremity of senescence phenotypes investigated, metric TT70 may be more informative than the duration of flag leaf senescence (EEtoLeafSen), which is more frequently used when recording senescence variation (Table 1). For example, in mild years plants may not reach terminal senescence, as we observed in 2016, preventing calculation of EEtoLeafSen. In contrast, TT70 scores can be calculated prior to harvest, increasing efficiency of phenotypic selection. On a cautionary note, whilst we identified metrics TT70 and MeanPed as most informative under our conditions, this may be germplasm dependent and not universally true. For example, Shi *et al*. (2016) observed *Triticum aestivum* cv. Wenmai and cv. Lankaoaizao senesce non-sequentially, whereby flag leaves senesced prior to 2^nd^ leaves, whilst peduncles of *GPC-1* RNAi lines remained green (Cantu *et al*., 2011).

### Functional or not? Understanding the Relationship between Grain Filling and Leaf Senescence

A positive relationship between senescence and grain fill duration is often assumed, with grain fill duration and grain weight highly correlated, *r* = 0.77, *P* < 0.026 (Dias and Lidon, 2009). During QTL analysis of the *Triticum aestivum* cv. Spark × Rialto double haploid population, Simmonds *et al*. (2014) identified a single QTL on chromosome 6A associated with green canopy duration, TGW and yield. Using NILs contrasting for Rialto and Spark 6A alleles, yield, and senescence traits co-segregated. The reported grain filling extension was associated with earlier flowering and delayed grain maturation, which occur 1 day earlier and ∼2 days later, Rialto and Spark alleles respectively, not green canopy duration per se (Simmonds *et al*., 2014).

Differences in grain filling may be environmentally dependent. For example, under glasshouse conditions no differences in grain maturation were recorded for *gpc-1* (*NAM-1*) mutants, despite senescence being delayed by 25 days (Simmonds *et al*., 2014; Borrill *et al*., 2015), whereas differences were observed amongst wheat lines carrying *NAM-A1* variants in dryland environments (Alhabbar *et al*., 2018). Conversely, the extended grain fill duration reported for Staygreen A and B, which encode mutations in *NAM-A1* and *NAM-D1* respectively, was consistent over multiple years (Chapman *et al*., 2020), contributing to the observed, heading-independent, grain maturity differences (Figure 7).

The extended photosynthetic duration associated with functional staygreen phenotypes may provide additional resources for grain filling, supporting trait deployment in breeding. Xie *et al*., (2016) hypothesise a delay in onset combined with a rapid rate of senescence maximises grain weight and yield potential, especially as grain filling rate (growing °C days) and grain weight are correlated, *r* = 0.91, *P* = 0.002 (Dias and Lidon, 2009). Grain filling rate can be indirectly assessed through scoring senescence progression. For example, Xie *et al*. (2016) found rates of maximum chlorophyll loss and average grain fill to be correlated, *r* = 0.27 to 0.35, *P* < 0.01, however correlations between senescence metrics and grain fill duration were inconsistent, *r* = −0.4 to 0.4, *P* < 0.01 to *>* 0.05. Together, this emphasises the need to score both time to terminal senescence and grain maturation, and would confirm the relationship assumed by Lopes and Reynolds (2012) and Pinto *et al*. (2016). We demonstrate subjecting grain to thumbnail impressions can sufficiently characterise differences in grain maturation (Figure 7), providing a means of rapid assessment. Alternatively, if an objective assessment of grain maturation is required we suggest recording spike moisture content from ∼6 weeks after anthesis, as conducted for *NAM-1* mutants (Avni *et al*., 2014).

Time to maximum senescence rate can coincide with maximal grain dimensions (Xie *et al*., 2015), therefore maintaining synchronicity of senescence and grain filling processes is of concern when identifying or selecting staygreen traits. As photosynthesis terminates halfway through the rapid grain filling phase any delay could reduce remobilisation efficiency, as this marks the point when translocation and remobilisation of stored reserves, fructose and sucrose occurs (Takahashi *et al*., 2001). Phenotyping of leaf and peduncle senescence, alongside grain maturity could deliver insights into the process, and lines displaying ‘green leaf and ripe ear’ phenotypes should be selected against (Figure 3F). Conversely, the grain fill extension reported for Staygreens A and B (Chapman *et al*., 2020) delayed grain maturation (Figure 7) which could disrupt harvest, or adversely affect wheat quality through altering deposition of triticin, glutenin and gliadin storage proteins (Takahashi, 2001), requiring further investigation.

### Variable Trait Expression: Consider Target Environments

Staygreen traits are associated with conveying tolerance to heat, drought and low nitrogen conditions (Gregersen *et al*., 2013; Jagadish *et al*., 2015), with trait expression strongly influenced by the environment. For example, of the 19 senescence QTLs mapped for the *Triticum aestivum* cv. Ventor × Karl92 RIL population, segregating for temperature responses, only 3 were environmentally stable (Vijayalakshmi *et al*., 2010). Individually these 3 QTLs explained between 10 to 51 % of senescence variation. Conversely, variation explained by the 7 and 9 senescence QTLs identified exclusively under high or optimal temperature conditions were lower, averaging R^2^ = 0.18 ± 0.11 and R^2^ = 0.14 ± 0.07 respectively (Vijayalakshmi *et al*., 2010). Therefore, screening for senescence traits under different environments may help identify, potentially advantageous, stress adaptive QTLs for use in breeding, alongside major stable genetic regulators.

Water limitation also influences senescence. Estimated heritability of senescence traits recorded for the *Triticum aestivum* cv. Reeder × cv. Canan RIL population reduced from H^2^ = 0.78 under irrigated conditions to H^2^ = 0.51 to 0.81 when rainfed (Naruoka *et al*., 2012). Between-year weather variation can help identify putative epistatic interactions, as documented during QTL mapping of the *Triticum aestivum* cv. Chirya-3 × Sonalika RIL population (Kumar *et al*., 2010). Kumar *et al*. (2010) identified an environmentally stable senescence QTL located on chromosome 1AS, which in combination with year-dependent QTLs located on 3BS (2005) and 7DS (2006) accounted for up to 38.7 % of staygreen trait variation. Whilst we identified both our staygreen traits as highly environmentally stable (Figure 5) we also recognise the influence of environment, as senescence was accelerated in 2017 and 2018 compared to 2016 due to increased temperature and water limitation.

Together, these examples illustrate the need for repeated, multi-environment, trialling to assess stress adaptivity or stability of senescence phenotypes. However, within breeding programmes lines are typically selected under high-input conditions, preventing phenotypic expression and selection of potential stress adaptive staygreen phenotypes. From our experience, multi-environment, multi-year trials allowed us to identify potential penalties associated with adoption of staygreen traits alongside appropriate target breeding environments. For example, although loss of glume colour and grain ripening occurred ahead of flag leaf senescence in 2016 (Figure 3F), supporting non-adoption of staygreen traits (Jenner *et al*., 1991; Barraclough *et al*., 2014), results of continental trials identified the trait as stress adaptive. Whilst certain ‘extreme’ staygreen phenotypes may be agronomically unsuitable, obtaining such knowledge helps identify environmental niches for which adoption of staygreen traits could provide maximum benefit.

## Conclusion

Improving our understanding of senescence requires adoption of a whole plant phenotyping approach. When mapping staygreen traits underpinned by *NAM-1* mutations, we found TT70 and MeanPed to be the most stable and discriminative metrics. Thumbnail impressions can effectively detect variation in grain maturity associated with senescence traits, providing a rapid means of assessment. In combination we hope these insights can help qualify senescence traits, and aid identification and selection of senescence variation for both breeders and researchers alike.

## Supplementary Material

**Supplemental Table 1** Year-Pairwise Senescence metric Spearman’s rank correlations for Staygreen A and Staygreen B RILs

## Acknowledgements

The authors wish to thank JIC and KWS field experimentation teams who make field trial plans a reality. Statistical analysis was performed following assistance from Luzie Wingen. Thanks go to Sophie Harrington, Phillipa Borrill and Chris Burt for encouraging E.A.C. to share experience of field phenotyping through writing this article, and to Gemma Molero for engaging with my ideas. Additional thanks go to Sophie Harrington for providing much-appreciated editorial assistance. The *Triticum aestivum* cv. Paragon EMS mutant population used was developed by Robert Koebner and Leodie Alibert.

*The authors declare that the research was conducted in the absence of any commercial or financial relationships that could be construed as a potential conflict of interest.*

