## Supplemental Table 1 for "Capturing and Selecting Senescence Variation in Wheat"

**Table 1.** Year-Pairwise Spearman’s rank correlations for senescence metrics recorded for Staygreen A and Staygreen B RILs. correlations calculated for the 36-42 RILs per population grown at Church Farm, Norwich, from 2016 to 2018, mean value per line, n = 3 (2017), n = 2 (2018). *P,* * < 0.05, ** < 0.01, *** < 0.001. Dash indicates correlation unable to be calculated as data not collected in 2016.

| **RIL Subset** | **Staygreen A** | | | **Staygreen B** | | |
| --- | --- | --- | --- | --- | --- | --- |
| **Year-Pair** | **2016 vs. 2017** | **2016 vs. 2018** | **2017 vs. 2017** | **2016 vs. 2017** | **2016 vs. 2018** | **2017 vs. 2017** |
| **MeanLeaf** | 0.76 *** | 0.65 *** | 0.83 *** | 0.39 * | 0.26 | 0.57 *** |
| **EEtoLeafSen** | -0.12 | 0.10 | 0.75 *** | 0.10 | 0.06 | 0.68 *** |
| **LeafSenDur** | 0.39 * | 0.39 * | 0.87 *** | -0.02 | 0.21 | 0.38 ** |
| **MeanPed** | - | - | 0.91 *** | - | - | 0.62 *** |
| **EEtoPedSen** | - | - | 0.85 *** | - | - | 0.63 *** |
| **PedSenDur** | - | - | 0.45 ** | - | - | 0.11 |
| **TT25** | 0.66 *** | 0.47 ** | 0.58 *** | 0.29 | 0.10 | 0.30 * |
| **TT30** | 0.68 *** | 0.87 *** | 0.64 *** | 0.33 * | 0.22 | 0.42 *** |
| **TT40** | 0.76 *** | 0.70 *** | 0.74 *** | 0.48 ** | 0.17 | 0.52 *** |
| **TT50** | 0.81 *** | 0.68 *** | 0.78 *** | 0.42 ** | 0.27 | 0.58 *** |
| **TT60** | 0.79 *** | 0.73 *** | 0.83 *** | 0.52 *** | 0.30 | 0.55 *** |
| **TT70** | 0.78 *** | 0.78 *** | 0.84 *** | 0.50 *** | 0.37 * | 0.59 *** |
| **TT75** | 0.71 *** | 0.71 *** | 0.79 *** | 0.47 ** | 0.35 * | 0.56 *** |
| **TT80** | - | - | 0.80 *** | - | - | 0.58 *** |
| $\bar{\boldsymbol{x}}$ **correlation (significant traits only)** | 0.703 | 0.643 | 0.748 | 0.444 | 0.325 | 0.537 |
| **Range** | 0.392 – 0.806 | 0.394 – 0.781 | 0.454 – 0.911 | 0.328 – 0.515 | 0.255 – 0.372 | 0.303 – 0.678 |
